# Brain Responsivity to Emotional Faces Differs in Alcoholic Men and Women

**DOI:** 10.1101/496166

**Authors:** Marlene Oscar-Berman, Susan Mosher Ruiz, Ksenija Marinkovic, Mary M. Valmas, Gordon J. Harris, Kayle S. Sawyer

**Affiliations:** Psychology Research Service, VA Boston Healthcare System, Boston, MA, USA; Department of Anatomy & Neurobiology, Boston University School of Medicine, Boston, MA. USA; Department of Radiology, Massachusetts General Hospital, Boston, MA, USA; Department of Psychology, San Diego State University, San Diego, CA, USA; Sawyer Scientific, LLC, Boston, MA, USA

**Keywords:** Alcoholism, reward, fMRI, gender, memory, emotion

## Abstract

Inclusion of women in alcoholism research has shown that gender differences contribute to unique profiles of cognitive, emotional, and neuropsychological dysfunction. We employed functional magnetic resonance imaging (fMRI) of abstinent long-term alcoholics (21 women [ALCw] and 21 men [ALCm]) and demographically-similar nonalcoholic controls (21 women [NCw] and 21 men [NCm]) to explore how gender and alcoholism interact to influence emotional processing and memory. Participants completed a delayed match-to-sample emotional face memory fMRI task. While the results corroborated reports implicating amygdalar, superior temporal, and cerebellar involvement in emotional processing overall, the alcoholic participants showed hypoactivation of the left intraparietal sulcus to encoding the identity of the emotional face stimuli. The nonalcoholic participants demonstrated more reliable gender differences in neural responses to encoding the identity of the emotional faces than did the alcoholic group, and widespread neural responses to these stimuli were more pronounced in the NCw than in the NCm. By comparison, gender differences among ALC participants were either smaller or in the opposite direction (higher brain activation in ALCm than ALCw). Specifically, Group by Gender interaction effects indicated stronger responses to emotional faces by ALCm than ALCw in the left superior frontal gyrus and the right inferior frontal sulcus, while NCw had stronger responses than NCm. However, this pattern was inconsistent throughout the brain, with results suggesting the reverse direction of gender effects in the hippocampus and anterior cingulate cortex. Together, these findings demonstrated that gender plays a significant role in the profile of functional brain abnormalities observed in alcoholism.

## Introduction

Alcoholism is a common and harmful ^1,2^ condition that has been associated with differences in the processing of emotion, memory, and face identity. The brain regions associated with the encoding of face identity and emotion have been established, and research is beginning to indicate how the activity of these regions varies in conjunction with chronic long-term alcohol use disorder (AUD). Memory and facial emotion recognition are adversely affected by AUD ^3–6^. Moreover, impairments in processing facial emotional expressions have been well characterized ^7–10^, and can endure after months of sobriety ^11,12^.

Functional MRI (fMRI) of facial emotion processing among healthy adults elicits activation in a widespread network of brain areas including the fusiform gyrus, lateral occipital gyrus, superior temporal sulcus, inferior frontal gyrus, insula, orbitofrontal cortex, basal ganglia, and the amygdala ^13–16^. In the present study, we explored face encoding in particular, which typically relies upon prefrontal cortex, amygdala, hippocampus, fusiform, and lateral parietal regions ^17–19^. Processing of emotional facial expressions and emotional identity appear to be partially functionally segregated ^20^; therefore, attention to identity of faces rather than to the emotions expressed on them influences the network utilized. Attention to face identity typically activates fusiform and inferior temporal areas, whereas attention to the emotional expression would more likely yield activation in superior temporal, amygdala, and orbitofrontal regions ^21–25^.

Functional neuroimaging studies of working memory for emotional content in AUD are relatively rare. One study that examined face memory encoding in a mixed-gender group of alcoholics found right lateralization of parahippocampal activation among controls, but not among alcoholic participants ^26^. A study examining explicit emotional face decoding in alcoholism (without a memory component) found that alcoholic men were not impaired at decoding facial emotional expressions, but used different brain regions to perform the task than did controls ^27^. In that study, participants rated the intensity of facial emotional expressions during functional MRI scans. The largest differences in blood oxygenation level-dependent (BOLD) responses were noted in the anterior cingulate cortex, with alcoholic men showing less activation than controls in this region in response to facial expressions of fear, disgust, and sadness. Another study that examined alcoholic men’s BOLD responses to emotional faces and words, wherein subjects made judgments about physical and perceived psychological attributes of facial images, reported a diminished amygdala response during deep encoding of faces ^28^.

Historically, AUD has afflicted more men than women, but prevalence for women has increased such that they are approaching similar rates ^5,29–32^. Much of what has been learned about the long-term effects of alcoholism has been based upon research that focused on men, or has not differentiated the results obtained from different genders. There exist clear differences in how alcohol affects men and women physiologically and in how they progress from social to problem drinkers ^31^. For example, we observed gender-dimorphic effects in multiple domains including emotional processing, personality, and drinking motives ^5,33^, brain white matter ^34–37^, morphometry of the brain reward system ^38^, and brain activity in response to emotional cues ^39^. However, there are few fMRI studies of gender-specific brain activation in alcoholism.

The present study sought to characterize abnormalities in neural activation among chronic long-term alcoholics through analysis of BOLD responses to photographs of faces that varied in emotional expressions. We were particularly interested in observing activation responses in brain regions that subserve face processing, memory encoding, and emotions, and to determine how these effects differ between alcoholic men (ALCm) and alcoholic women (ALCw), as compared to nonalcoholic men (NCm) and nonalcoholic women (NCw).

## Methods

### Participants

Participants included 42 abstinent long-term alcoholics (21 ALCw) and 42 nonalcoholic controls (21 NCw) (Supplemental Table 1). Participants were recruited within the Boston area through advertisements placed in public places (*e.g.*, hospitals, churches, stores), newspapers, and internet websites. The study was approved by Institutional Review Boards at our affiliated institutions. Participants gave written informed consent prior to participation, and were compensated for their time.

Selection procedures included an initial structured telephone interview to determine age, level of education, health history, and history of alcohol and drug use. Included participants were right-handed, had normal or corrected-to-normal vision, and spoke English as a first language. Participants were interviewed further to determine use of alcohol and other drugs, including prescription drugs that would affect the central nervous system. Current drug use excepting nicotine and caffeine was cause for exclusion. Criteria for exclusion also included history of liver disease, epilepsy, head trauma resulting in loss of consciousness for 15 minutes or more, HIV, symptoms of Korsakoff’s Syndrome or dementia, and schizophrenia. Additionally, individuals who failed MRI screening (e.g., metal implants and obesity) were excluded.

Neuropsychological testing was conducted at the Department of Veterans Affairs (VA) Boston Healthcare System facility. Participants completed a medical history interview, vision test, handedness questionnaire ^40^, Hamilton Rating Scale for Depression ^41^ and the Diagnostic and Statistical Manual (DSM-IV) Diagnostic Interview Schedule ^42^. Participants also were administered the Wechsler Adult Intelligence Scale (WAIS-III or WAIS-IV) and the Wechsler Memory Scale (WMS-III or WMS-IV) ^43,44^.

Alcoholic participants met DSM-IV criteria for alcohol abuse or dependence, and consumed 21 or more alcoholic drinks per week for five or more years. The extent of alcohol use was assessed by calculating Quantity Frequency Index (QFI) scores ^45^. QFI approximates the number of drinks consumed per day, and the amount, type, and frequency of alcohol consumption either over the last six months (control participants), or over the six months preceding cessation of drinking (alcoholic participants) to yield an estimate of ounces of ethanol per day. Alcoholic participants were abstinent for at least four weeks before the scan date. Control participants who had consumed 15 or more drinks per week for any length of time, including binge drinking, were excluded.

### Functional Imaging Task

Participants were given a delayed match-to-sample memory task in an MRI scanner, whereby they were asked to encode two faces that had one of three emotional valences: positive, negative, or neutral (Supplemental Figure 1). The face stimuli were shown in grayscale and were taken from a set of faces used in a previous study ^28^. Face stimuli were displayed simultaneously for three seconds, followed by an asterisk (*) for one second. Participants were asked to maintain these faces in memory while distractor stimuli were shown (for three seconds), immediately followed by a probe face (shown for two seconds) to assess memory for face identity, and ending with a variable-length fixation (two to thirty seconds, average ten seconds). Results obtained from analyses of the distractor and probe conditions are part of a separate research project. The participants responded to the probe face with their right index and middle finger, and psychophysiological recordings were taken from the left hand ^46^. The event-related design used nine six-minute runs with 18 trials per run, for a total of 162 trials. There were 54 trials for each emotional face valence. Counter-balancing and inter-trial intervals were calculated with optseq2 ^47^.

### Image Acquisition

Imaging was conducted at the Massachusetts General Hospital, Charlestown, MA. Data were acquired on a 3 Tesla Siemens (Erlangen, Germany) MAGNETOM Trio Tim MRI scanner with a 12-channel head coil. Sagittal T1-weighted MP-RAGE scans (TR = 2530 ms, TE = 3.39 ms, flip angle = 7°, FOV = 256 mm, slice thickness = 1.33 mm, slices = 128, matrix = 256 × 192) were collected for all subjects. Echo planar fMRI scans were acquired axially with 5 mm slice thickness and 3.125 × 3.125 mm in-plane resolution (64 × 64 matrix), allowing for whole brain coverage (32 interleaved slices, TR = 2 s, TE = 30 ms, flip angle = 90°). Within each six-minute run, 180 T2*-weighted volumes were obtained. Functional volumes were auto-aligned to the anterior/posterior commissure line to ensure a similar slice prescription was employed across participants. Prospective Acquisition Correction (3D-PACE) was applied during acquisition of the functional volumes to minimize the influence of participants’ body motion ^48^. A laptop running Presentation version 11.2 software (NeuroBehavioral Systems, Albany, CA) was used for visual presentation of the experimental stimuli and collection of participants’ responses. Stimuli were back-projected onto a screen at the back of the scanner bore and viewed by participants through a mirror mounted to the head coil. Participants wore earplugs to attenuate scanner noise.

### Structural Image Processing

Cortical surfaces were reconstructed using Freesurfer version 4.5.0 (http://surfer.nmr.mgh.harvard.edu) to obtain segmentation labels ^49,50^ along with white matter and exterior cortical surfaces ^51^. These were visually inspected on each coronal slice for every subject, and manual interventions (*e.g.*, white matter volume corrections) were made as needed. The Destrieux atlas parcellation for FreeSurfer ^52^ and subcortical segmentation were used to define anatomical regions of interest (ROI) in the functional analyses.

### Functional Image Processing and Statistical Analyses

Effects of Group, Gender, and Emotion on BOLD signal were evaluated using a whole-brain cluster analysis, as well as a ROI analysis. Processing of the functional data was performed using FreeSurfer Functional Analysis Stream (FS-FAST) version 5.3 (http://surfer.nmr.mgh.harvard.edu/), SPSS Version 17.0 (IBM, Chicago, IL, USA), and custom Matlab scripts (The MathWorks, Natick, MA).

Preprocessing of functional images for first-level (individual subject) analyses included motion correction, intensity normalization, and spatial smoothing with a 5-mm Gaussian convolution kernel at full-width half-maximum. BOLD response was estimated using a Finite Impulse Response (FIR) model, which allows for estimation of the time course of activity (percent signal change for a given condition) within a voxel, vertex, or ROI for the entire trial period. For each condition, estimates of signal intensity were calculated for 2 pre-trial and 10 post-trial onset TRs, for a total analysis window of 24 seconds. Motion correction parameters calculated during alignment of the functional images were entered into the analysis as external regressors. Alignment of the T2*-weighted functional images with T1-weighted structural volumes was accomplished through an automated boundary-based registration procedure ^53^. These automated alignments were manually inspected to ensure accuracy.

Statistical maps were generated for each individual subject for contrasts between experimental conditions. Contrasts for the facial emotion conditions included: (1) positive faces vs. fixation, (2) negative faces vs. fixation, (3) neutral faces vs. fixation, (4) positive faces vs. negative faces, (5) positive faces vs. neutral faces, and (6) negative faces vs. neutral faces. Analyses of each of these contrasts included removal of prestimulus differences between the contrasted conditions by averaging the first three time points (two pre-trial and one post-trial onset) for each condition and subtracting this mean from each time point for that condition. Time points summed for inclusion in each contrast were chosen to reflect peak stimulus-related activity: FIR estimates of hemodynamic responses to the emotion effects were examined during the time period of 2-10 seconds post emotional face onset.

Second-level (group) analyses on cortical regions were accomplished using a surface-based morphing procedure for intersubject alignment and statistics ^54^. Group-averaged signal intensities during each experimental condition relative to fixation were calculated using the general linear model in spherical space for cortical regions, and were mapped onto the canonical cortical surface *fsaverage*, generating group-level weighted random-effects *t*-statistic maps; the same procedure was performed for the volume with a subcortical mask. A 5 mm smoothing kernel (FWHM) was employed for all group and intergroup maps.

Intergroup comparison *t*-statistic maps were generated to examine between-group effects by contrasting: (1) alcoholic participants vs. control participants, (2) ALCm vs. NCm, (3) ALCw vs. NCw, (4) ALCm vs. ALCw, (5) NCm vs. NCw, (6) men vs. women. Additionally, Group by Gender interaction maps for each contrast were calculated.

For the cortical surface maps, a Cluster-level correction for multiple comparisons were applied to each map ^55^, wherein each vertex (or voxel) was required to meet two criteria to be considered significant: the *p*-value for the within-group or between-groups contrast *t*-test must be significant at *p* < 0.001, and the vertex (or voxel) must be contiguous with a set of vertices that span a minimum surface area of 100 mm^2^ (or voxels with a minimum volume of 300 mm^3^). Cortical surface cluster regions were identified by the location of each cluster’s peak vertex on the cortical surface according to the Destrieux atlas. Subcortical cluster regions were identified by the Talairach ^56^ coordinates of each cluster’s peak voxel according to the Talairach Daemon ^57^.

Anatomically-defined ROI were selected for the emotional faces BOLD analyses to include regions identified *a priori* that are known to be involved in the recognition of emotions, and in visual processing and memory encoding of human faces. These were the amygdala, fusiform gyrus, hippocampus, parahippocampal gyrus, intraparietal sulcus, orbitofrontal cortex, superior temporal gyrus and superior temporal sulcus. Left and right hemisphere regions were analyzed separately.

Statistical preprocessing and time course visualization of ROI data were performed using custom scripts written for Matlab version 7.4.0. Signal intensity for each region was averaged across all vertices (or voxels) included in the region for each condition on the individual participant level. To compute percent signal change for each participant within an ROI, signal estimate per condition and time point was divided by the average baseline activity for that participant in the same manner as for the statistical maps. Group and Group-by-Gender averages of the normalized time courses were computed for each condition, and were visualized by plotting the percent signal change for each condition at each time point (*i.e.*, TR) of the trial.

For the ROI analyses, percent signal changes of the BOLD signal within each ROI were entered as dependent variables into repeated-measures ANOVA models with between-group factors of Group (alcoholic or control) and Gender (men or women) and within-subjects factor of facial Emotion (positive, negative, or neutral).

## Results

### Participant Characteristics

Supplemental Table 1 summarizes means, standard deviations, and ranges of participant demographics, IQ and memory test scores, and drinking variables. The alcoholics and controls did not differ significantly by age (mean ages 54 years), and although NCw were older than NCm, control groups did not differ significantly from the respective alcoholic groups of the same gender. ALCm had one year less education than NCm. Groups did not differ significantly on WAIS-III Full Scale IQ scores. ALCw had higher Hamilton Depression Scale scores than ALCm. Alcoholic participants drank 11 drinks per day on average, had a mean duration of 17 years of heavy drinking, and were sober for an average of eight years.

### Neuroimaging Cluster Analyses

First, group-level cluster analyses of each facial emotion condition vs. fixation yielded clusters too large to be described in an anatomically-relevant way with a single peak location. Therefore, these data are summarized qualitatively (in the text) and illustrated with Figure 1. Next, group-level clusters are reported (Table 1) for emotion contrasts (i.e., positive vs. negative, positive vs. neutral, and negative vs. neutral). Intergroup clusters are reported for each emotion condition vs. fixation (Table 2), followed by the intergroup clusters for emotion contrasts, (described in the text).

**Table 1.**
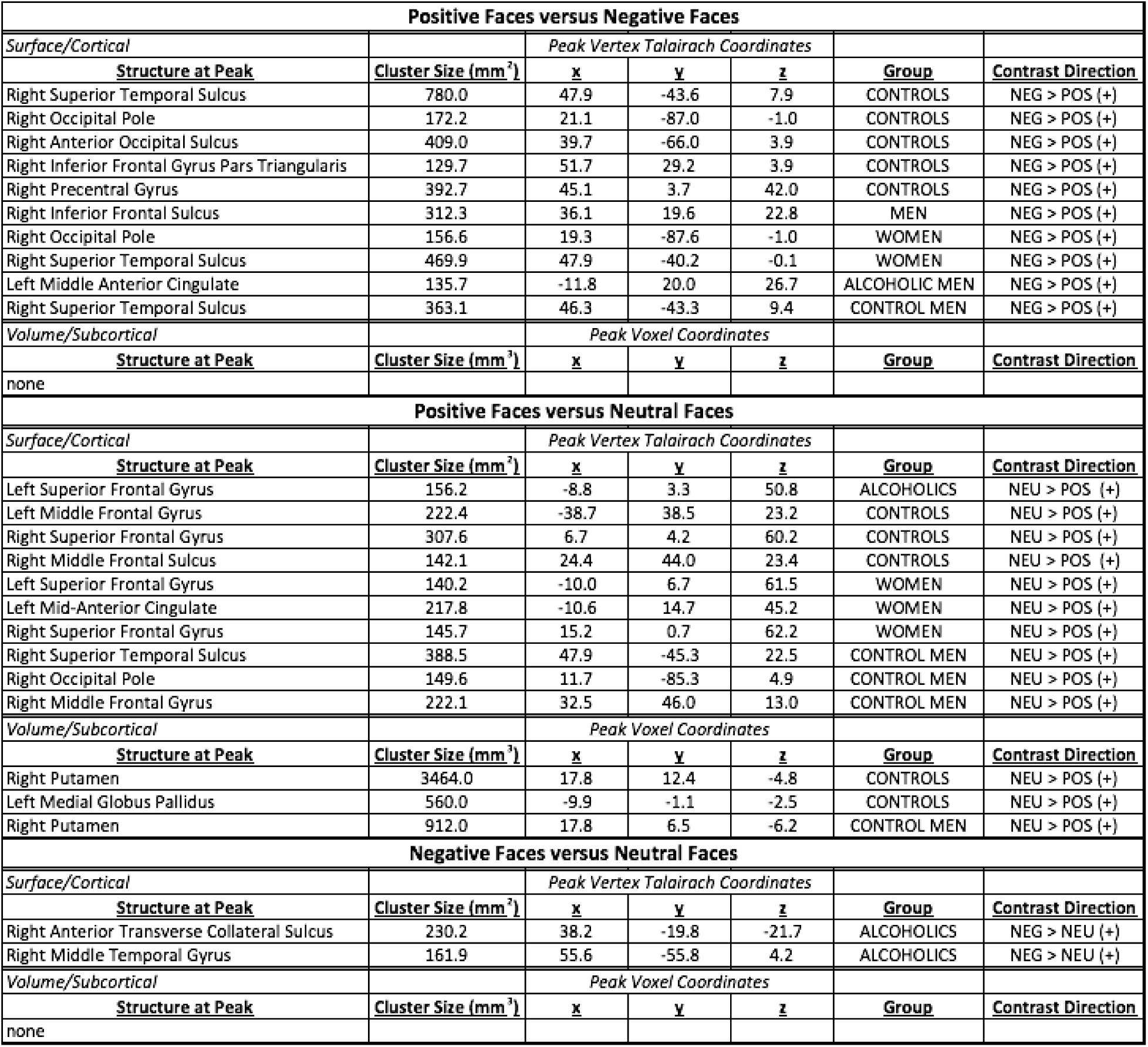
Emotion Whole Brain Group Cluster Analysis: Positive vs. Negative, Positive vs. Neutral, Negative vs. Neutral. Minimum groupwise significance for all vertices/voxels within a cluster set at p = 0.001. Cortical clusters were minimum 100 mm^2^. Subcortical clusters were minimum 300 mm^3^. Contrast directions are reported as absolute values of effects for each condition vs. fixation and denoted as (+) for increases relative to fixation and (-) for decreases relative to fixation.

**Table 2.**
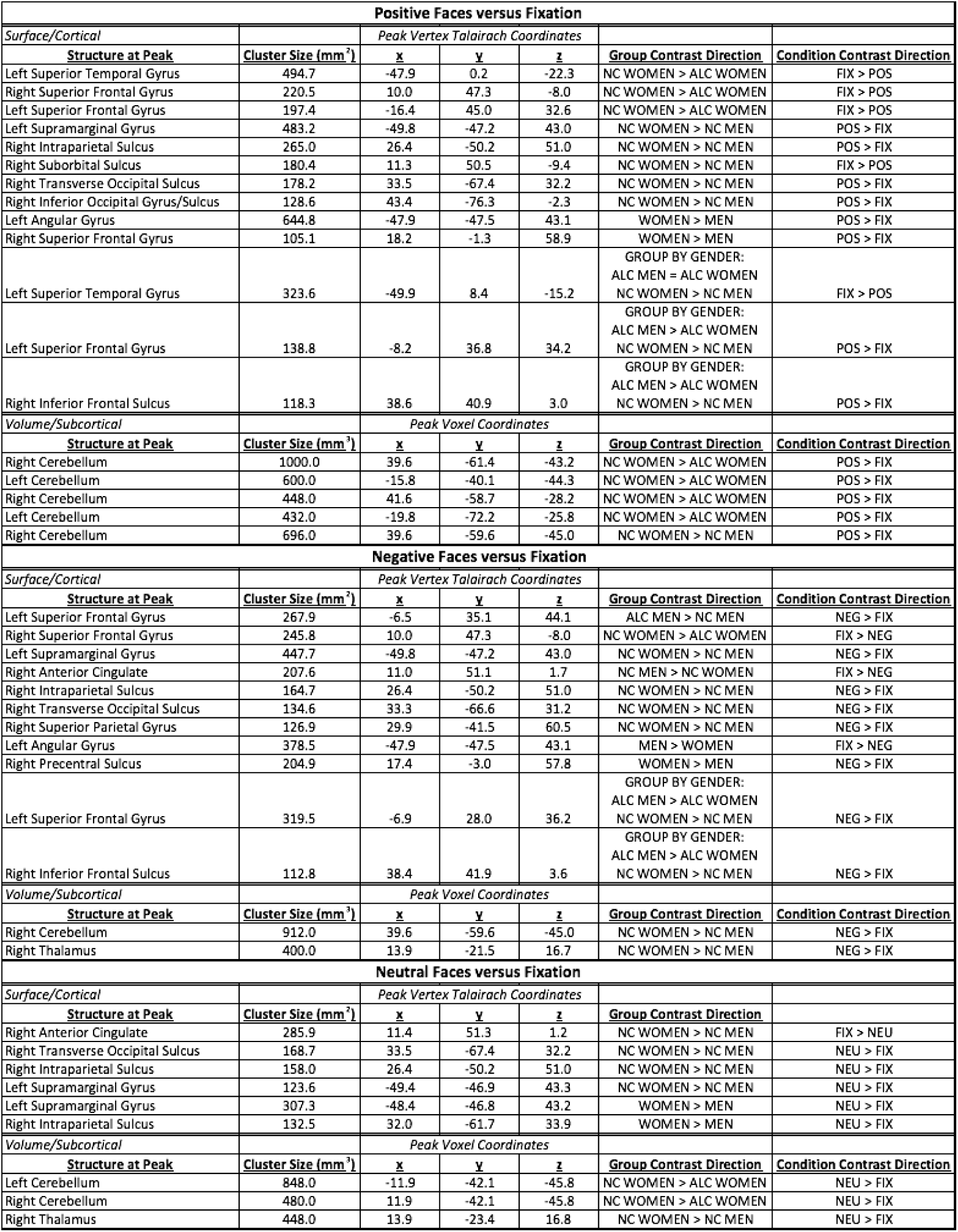
Emotion Whole Brain Intergroup Cluster Analysis II: Positive vs. Fixation, Negative vs. Fixation, Neutral vs. Fixation. Minimum between-group significance for all vertices/voxels within a cluster set at *p* = 0.001. Cortical clusters were minimum 100 mm^2^. Subcortical clusters were minimum 300 mm^3^.

Alcoholics and controls of both genders utilized a distributed network of cortical brain regions to process faces of all three emotional valences as compared to the fixation stimulus. As an example, Figure 1 shows the t-statistic cluster map of the contrast of positive vs. fixation displayed on the lateral surface; the medial and lateral views of the negative vs. fixation and neutral vs. fixation are shown in Supplemental Figures 2–6. Subcortical volume-based t-statistic cluster maps are shown in Supplemental Figures 7–10. Several *face-activated* regions were more responsive when participants viewed the face stimuli than during the fixation condition: dorsolateral prefrontal cortex, motor cortex, anterior insula, inferior temporal cortex (including fusiform), parietal cortex, the occipital lobes, limbic structures, basal ganglia, and the cerebellum. A different set of *fixation-activated* regions was more active during fixation than during the face conditions, forming the network known as the default mode network, because those regions typically are more active during rest than during attentionally-demanding cognitive tasks (Buckner et al., 2008; Cavanna and Trimble, 2006; Gusnard et al., 2001; Kim and Lee, 2011). The regions making up this network include the superior frontal cortex, medial prefrontal cortex, medial temporal lobe structures, the middle temporal gyrus, the posterior cingulate cortex plus precuneus, and the angular gyrus.

**Figure 1.**
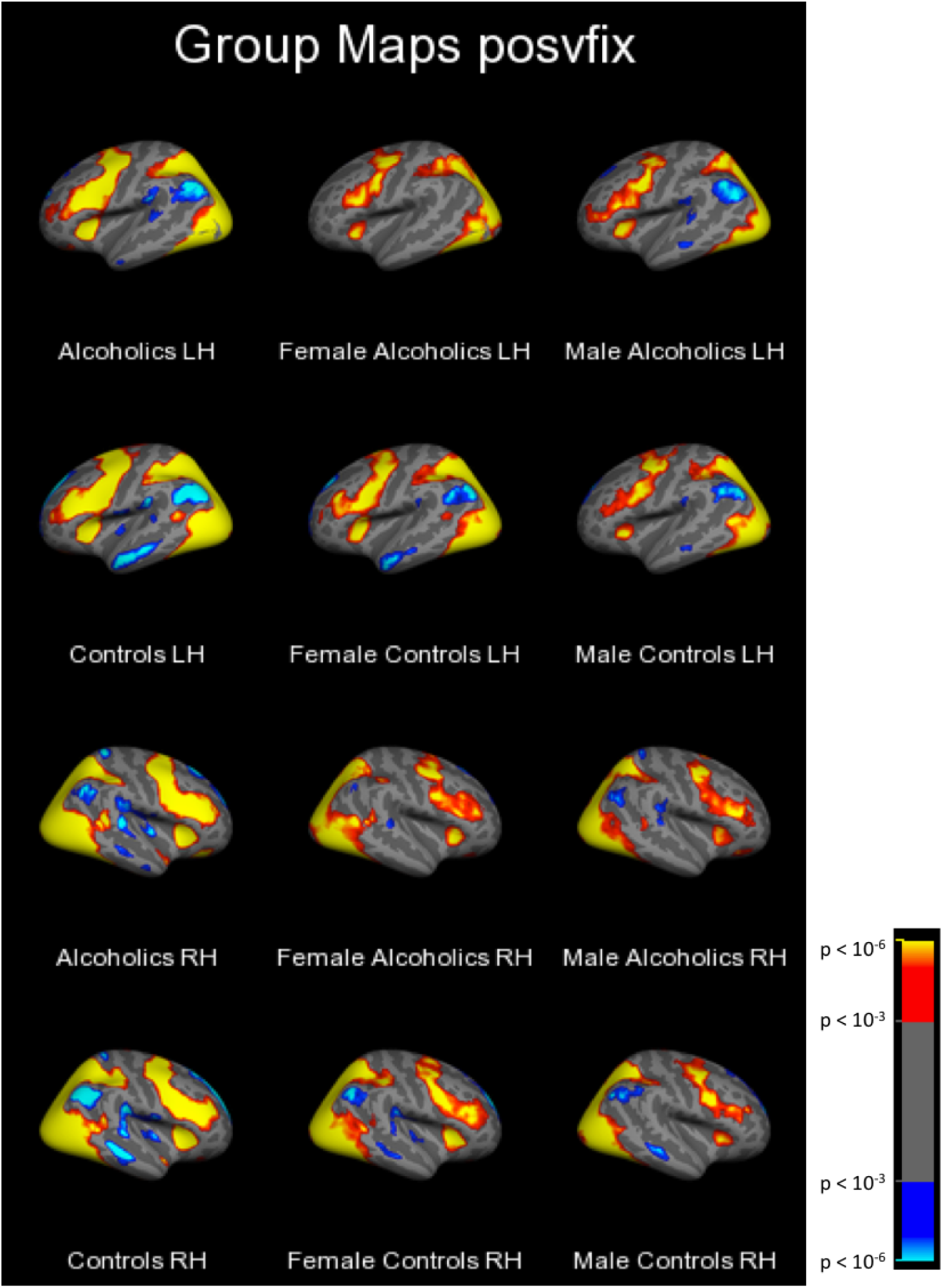
Group Cortical Surface Cluster Maps: Positive vs. Fixation, Lateral. Cluster-corrected at *p*<.001 with minimum cluster size 100 mm^2^.

As was observed for the contrast of faces vs. fixation, results for the emotion contrasts revealed several regions, but the clusters were of smaller spatial extent and less consistent across participant groups. Significant clusters are summarized in Table 1. Positive faces elicited the least activation, as compared to both neutral and negative faces. These effects were significant primarily in regions of the frontal lobes, although additional regions identified were in the occipital and temporal lobes, along with basal ganglia structures.

Clusters of between-group differences on each emotion condition vs. fixation are described in Table 2. The results were complicated and differed by brain region, contrast direction, and group comparison. For positive faces, significant clusters were identified within all lobes of the brain, and in the cerebellum. The group contrast directions for all of these clusters indicated greater activation values in the subgroup of NCw compared to the ALCw and NCm, and the effects were observed across both fixation-activated (default mode) and face-activated regions. Significant Group by Gender interaction effects for temporal and frontal regions were driven by the lower activation of NCm than NCw, while ALCm had similar or greater activation than ALCw. The negative faces revealed a pattern of group differences that encompassed many brain regions. For parietal and occipital regions, NCw had greater values than NCm, but the reverse (NCm > NCw) was observed for the anterior cingulate. As was found for the positive faces, two frontal clusters were identified where significant Group by Gender interactions were driven by lower negative vs. fixation contrast values obtained from NCm than NCw and higher contrasts from ALCm than ALCw. The neutral faces revealed many clusters as well, with NCw having greater contrasts than NCm in clusters across the parietal lobe, cerebellum, and limbic structures.

The three clusters that were identified with significant group differences for contrasts between emotional face conditions had peak voxels contained within the left and right amygdala (872 mm^3^ and 384 mm^3^, respectively), and left hypothalamus (464 mm^3^). For the clusters with peak voxels located in the amygdala, NCm had greater activity than NCw in which the contrast showed greater activation for positive than negative faces. For the cluster that included the left hypothalamus, men showed greater activation than women (combined across Group) for the contrast identifying regions where neutral faces had greater activation than positive faces.

### Neuroimaging Region of Interest Analyses

Results of repeated-measures ANOVAs examining between-subjects effects of Group and Gender and within-subjects effects of facial Emotion on BOLD percent signal change within each ROI are reported below. Means and standard deviations represent the percent signal change across each ROI for the time period of two to 10 seconds post-face stimulus onset.

#### Intraparietal Sulcus

A significant main effect of Group was found for left intraparietal sulcus activation during encoding of the emotional faces (*F* = 4.172, *p* = 0.044) (Figure 2). Controls had more activity in this region during face encoding (*M* = 0.268, *SD* = 0.123) than did alcoholics (*M* = 0.212, *SD* = 0.130). Activity in the right intraparietal sulcus did not vary significantly by Group, Gender, or facial Emotion.

**Figure 2.**
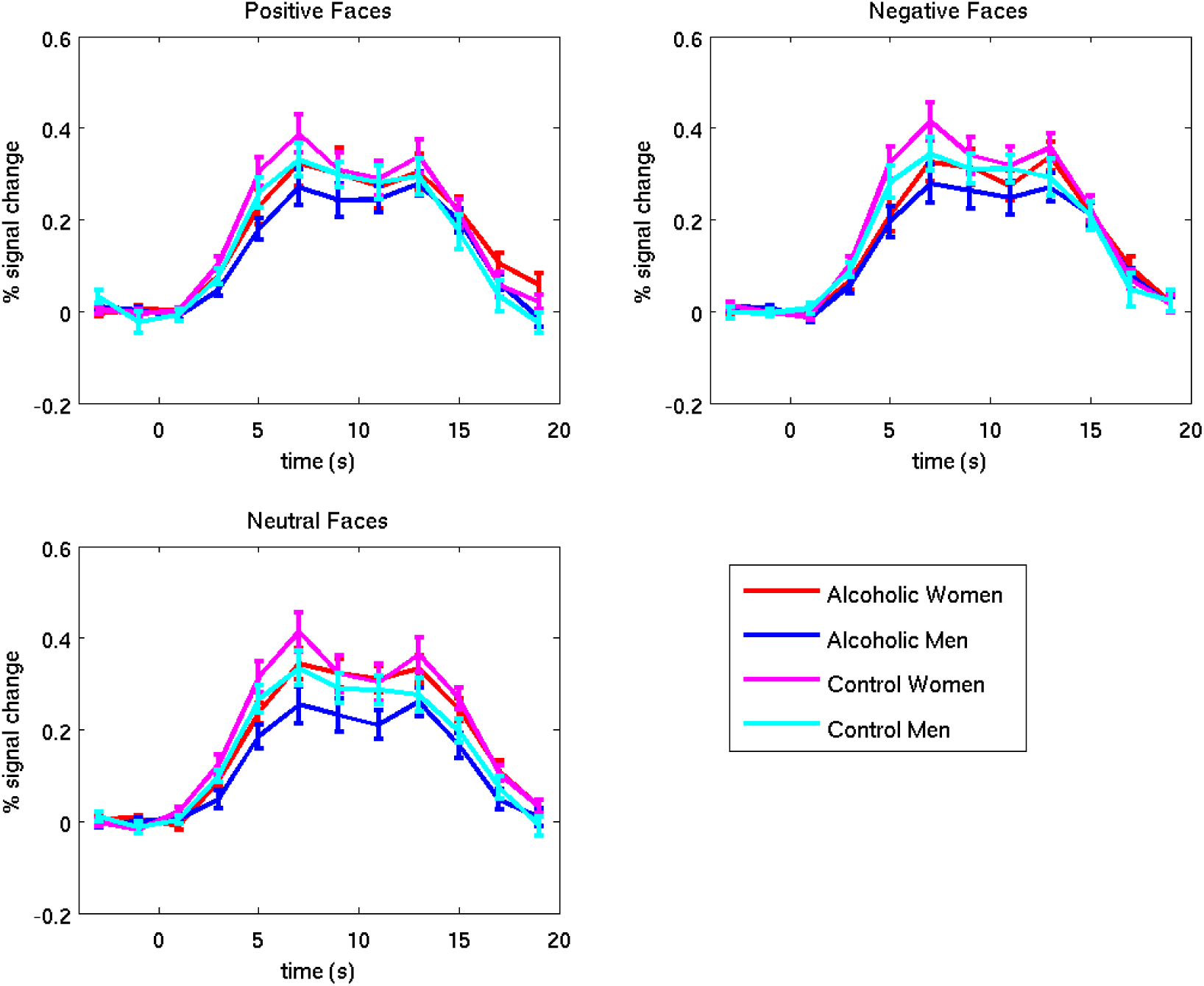
Left Intraparietal Sulcus. Error bars represent SEM. The two peaks represent brain activity resulting from the encoded faces and distractor. The analysis window used to examine the faces was 2 to 10 seconds.

#### Hippocampus

A significant Emotion by Group by Gender interaction was identified (*F_1,80_* = 4.005, *p* = 0.049), wherein NCm showed a trend toward a main effect of Emotion (*F_1,20_* = 3.297, *p* = 0.084), but this was not found in NCw (*F_1,20_* = 2.387, *p* = 0.352), ALCm (*F_1,20_* = 0.013, *p* = 0.909), nor ALCw (*F_1,20_* = 0.727, *p* = 0.404) (Figure 3). NCm tended toward more activation in the left hippocampus in response to positive faces (*M* = 0.044, *SD* = 0.087) relative to neutral faces (*M* = 0.011, *SD* = 0.105) (*t_20_* = 1.816, *p* = 0.084), but there was no difference between positive and negative (*M* = 0.022, *SD* = 0.073) faces (*t_20_* = 1.570, *p* = 0.132), nor between negative and neutral faces (*t_20_* = 0.834, *p* = 0.414). A trend for a Group by Gender effect (*F_1,80_* = 3.286, *p* = 0.074) was found in left hippocampus activation in response to faces, with ALCw (*M* = 0.022, *SD* = 0.103) having higher levels of activity in this region than ALCm (*M* = −0.009, *SD* = 0.103), and with NCm (*M* = 0.025, *SD* = 0.103) having higher levels of activity than NCw (*M* = −0.002, *SD* = 0.103). Activity in the right hippocampus did not vary significantly by Group, Gender, or facial Emotion.

**Figure 3.**
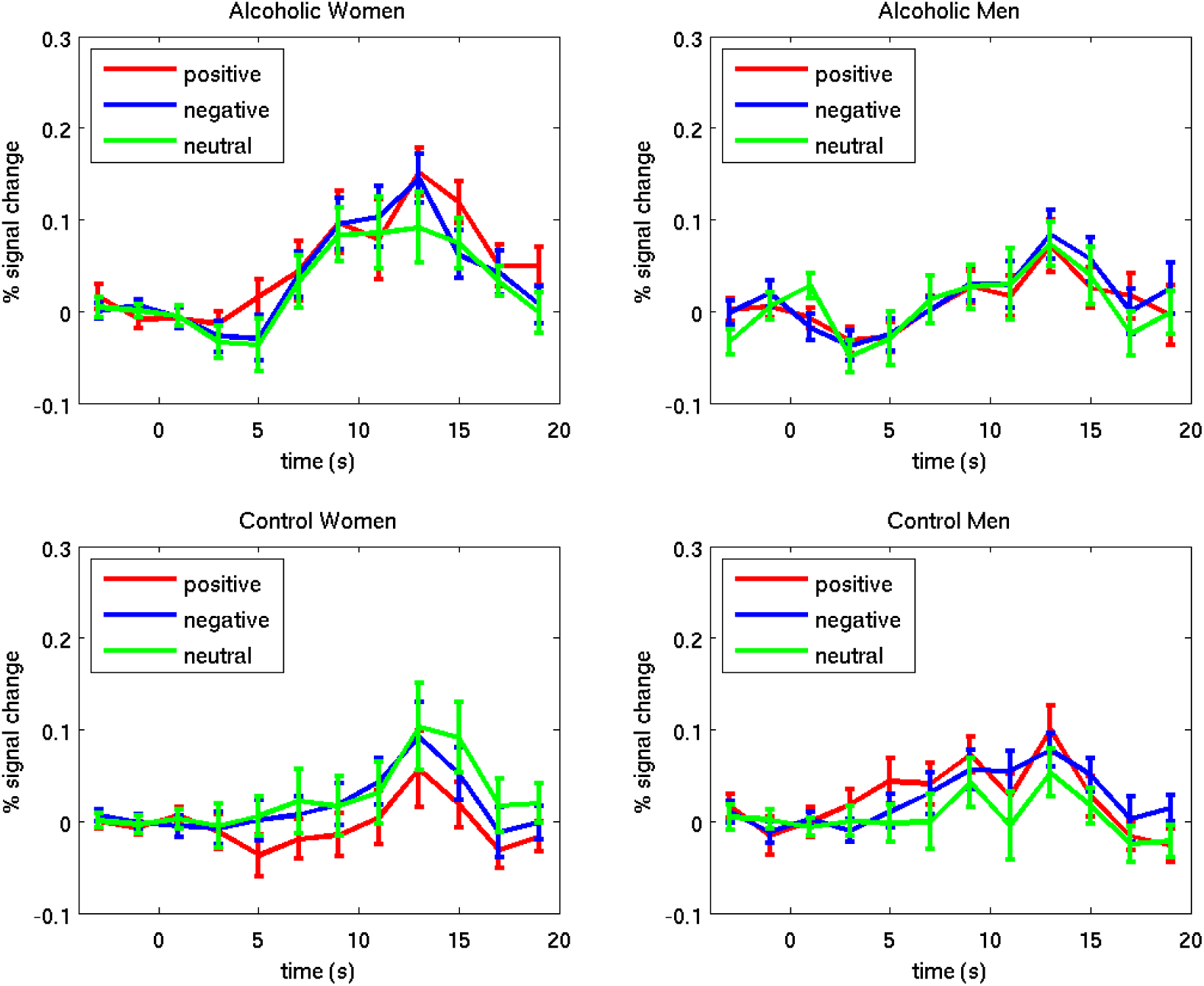
Left Hippocampus. Error bars represent SEM. The percent signal change represents brain activity resulting from the encoded faces and distractor. The analysis window used to examine the faces was 2 to 10 seconds.

#### Amygdala, fusiform, orbitofrontal cortex, parahippocampal cortex, superior temporal gyrus, and superior temporal sulcus

Activity in these regions did not vary significantly by Group, Gender, or facial Emotion.

In summary, for region of interest analyses, we observed higher responsivity during face encoding among controls than among alcoholics in the left intraparietal sulcus, a region that has been identified as playing an important role in focusing attention to enhance working memory ^58^. In the left hippocampus, an important memory structure, a significant interaction of Emotion, Group, and Gender, indicated that (a) the NCm activated this region more to positive faces than to neutral faces compared to the other subgroups, and (b) the ALCw activated more to the faces irrespective of valence, compared to the other subgroups.

## Discussion

In the present study, whole-brain group-level cluster analyses comparing activation among facial emotion conditions showed that participants had higher responses to negative than to positive faces in right hemisphere superior temporal sulcus, occipital, and inferior frontal regions. Consistent with our results, the superior temporal sulcus has been implicated in differentiating emotional facial expressions ^20,21,59^. Increased inferior frontal activation in response to negative relative to positive faces has been shown during emotional face recognition ^60^. Subcortically, among controls, neutral faces elicited greater activation than positive faces in the putamen and pallidum. A meta-analysis of emotional face processing included the putamen as a key structure ^61^. Additionally, the putamen was shown to respond to neutral faces in a functional neuroimaging study that used neutral faces to assess effects of attractiveness ^62^. The alcoholic group’s responses to negative faces were greater than to neutral faces in the collateral sulcus (adjacent to the anterior fusiform gyrus). While the fusiform is more commonly implicated in processing identity information ^20^, some studies also have implicated the collateral sulcus in processing emotional facial expressions ^21,59^.

Examination of whole-brain intergroup cluster contrasts revealed a gender difference among controls wherein the difference in responses to positive and negative faces was larger among NCm than NCw in the amygdala, a region with an undisputed role in emotional processing ^63,64^. The cerebellum also has been implicated in the processing of emotions ^65,66^, and emotional faces in particular ^61,67^. In the present study, NCw had significantly higher responses to both positive and neutral faces in the cerebellum than did ALCw, indicating abnormal function in addition to previous reports of abnormal cerebellar structure in relation to alcoholism ^68,69^. A region of the superior frontal gyrus extending to the anterior cingulate was identified wherein ALCw had less fixation-related activation as compared to positive faces than did NCw. This result mimics a finding among social phobics ^70^, indicating higher default mode network activation to fixation compared to activation during emotional face perception.

Interactions of group and gender were identified in responses to negative faces in the left superior frontal gyrus and the right inferior frontal sulcus. In both these regions, ALCm had higher responses to negative faces than did ALCw, whereas among controls, women had higher responses than men. ALCm also had higher responses to negative faces than did NCm in the left superior frontal gyrus. Taken together, these findings are suggestive of hyper-responsivity among ALCm to negatively-valenced faces.

Region of interest analyses examining effects of Group, Gender, and Emotional face valence yielded few significant results. In the left intraparietal sulcus, a region which has been singularly identified as playing an important role in focusing attention to enhance working memory ^58^, we found higher responses during face encoding among controls than among alcoholics. An interaction of Emotion, Group, and Gender was found in the left hippocampus, showing that only NCm activated this region more for positive faces than for neutral faces.

The amygdala, superior temporal sulcus, and orbitofrontal cortex are commonly reported to be sensitive to emotional face valence ^17,20,61^, but our results could not indicate whether the effect was in the positive or negative direction, only that the results of the ROI analyses were too small to detect with our paradigm and sample. There may have been several reasons for the modest findings in these areas. ROI were chosen based on neuroanatomically defined areas known to be implicated in emotional processing. Some of these regions, the superior temporal sulcus in particular, are relatively large, and emotion-related processing occurs only within a fraction of the region ^21^. Our analyses compared mean responses across an entire region, without the use of a functional mask. As such, strong but focal effects (as can be seen in the vertex-wise whole brain group map, Figure 1) between emotions within a region may have been diminished when averaged with other sections of that region where null or even diametrically opposing effects were found.

Another potential reason that large differences in emotional-related ROI were not observed could be that while two-thirds of the encoded faces displayed emotional expressions (*i.e.*, the positive and negative faces), the emotional expression of the faces were not explicitly relevant to the task, as it was the identity of the faces that was to be recalled. A number of studies have shown that activation of limbic regions, and particularly the amygdala, in response to emotional faces is dependent on the task-relevance of the emotional content of the faces ^71,72^. Another study examining memory for emotional face valence contrasted with face identity memory demonstrated higher orbitofrontal cortex activity when the emotional valence of the face was to be remembered relative to when the identity of the face was to be remembered ^17^. This study indicated that a single network of regions including the hippocampus, amygdala, and orbitofrontal cortex maintains both facial identity and facial emotional expressions in working memory during a delay period ^17^. The present study may have been more likely to find effects of emotion in these regions were the emotional valence the item to be remembered rather than the identity. Further, the study by LoPresti and colleagues (2008) found differences in orbitofrontal cortex activity between positive and negative faces in the explicit emotion memory task at the time of recall, but not at the time of encoding. Additionally, in a recent fMRI study employing stimuli eliciting stronger emotional reactions than facial expressions, we reported clear gender differences among alcoholics ^39^.

In summary, we have corroborated previous reports implicating amygdalar, superior temporal, and cerebellar involvement in emotional processing, and have further demonstrated more credible gender differences in neural responses to emotional faces among controls than among alcoholics. The findings from the present study can be viewed in the context of the Extended Reward and Oversight System (EROS) ^38,73^. EROS is comprised of interconnected cortical and subcortical regions, including prefrontal and cingulate cortices, the thalamus and hypothalamus, the nucleus accumbens, insula, and limbic structures including the amygdala and hippocampus. These regions are important for reinforcement of behavioral responses, primarily through modulation by cortical areas involved in memory, emotion, judgment, and decision-making. Importantly, components of EROS are structurally abnormal in alcoholic men and women ^38,73^.

### Limitations

Our results are based upon observational cross-sectional data, and as such, it is not possible to determine if chronic alcohol usage caused, or resulted from, the observed dysregulated emotional reactivity, or perhaps a combination of both. Further, we had limited information about the potentially confounding variable of smoking status, and therefore, it was not included in the analyses. Smoking abstinence has been associated with increased emotional reactivity in response to negative stimuli ^74^ and interactions with alcoholism ^75,76^, and therefore, may have influenced the results of the present study. The analyses of these data were performed in the context of a task that included the influence of intervening distractor and probe face images. Therefore, it is possible that interference from these images obscured differences that might otherwise have been observed. Future investigations of the interaction between emotional face valence and distractor conditions could resolve these effects. Despite these considerations, the present findings highlight the need for continued research on the overlap between gender differences in processing of emotional stimuli and the development of pathological alcohol consumption.

## Acknowledgements

This work was supported by funds from the US Department of Veterans Affairs Clinical Science Research and Development grant I01CX000326; the National Institute on Alcohol Abuse and Alcoholism (NIAAA) of the National Institutes of Health, US Department of Health and Human Services, under Award Numbers R01AA07112, R01AA016624, K05AA00219, and K01AA13402; and shared instrumentation grants 1S10RR023401, 1S10RR019307, and 1S10RR023043 from the National Center for Research Resources (now National Center for Advancing Translational Sciences) at the Athinoula A. Martinos Center, Massachusetts General Hospital. The authors thank Elinor Artsy, Sheeva Azma, Anne-Mette Guldberg, Zoe Gravitz, Doug Greve, Steve Lehar, Diane Merritt, Alan Poey, Elizabeth Rickenbacher, Trinity Urban, Jennifer Wilkens, and Robert Zondervan for assistance with manuscript preparation and recruitment, assessment, analysis, or neuroimaging of the research participants. The content is solely the responsibility of the authors and does not necessarily represent the official views of the National Institutes of Health, the U.S. Department of Veterans Affairs, or the United States Government.

## Supplemental Material

**Supplemental Table 1.** Participant characteristics.

|  | ALCOHOLICS |  |  | CONTROLS |  |  |
| --- | --- | --- | --- | --- | --- | --- |
|  | All<br>*n* = 42 | Women<br>*n* = 21 | Men<br>*n* = 21 | All<br>*n* = 42 | Women<br>*n* = 21 | Men<br>*n* = 21 |
| **Age<sup>a</sup> (years)** |  |  |  |  |  |  |
| *mean* | 53.9 | 53.4 | 54.4 | 53.9 | 57.7 | 50.2 |
| *standard deviation* | 11.0 | 11.4 | 10.8 | 12.4 | 13.6 | 10.1 |
| *range* | 26.5 - 76.7 | 26.5 - 73.0 | 26.6 - 76.7 | 25.8 - 76.9 | 25.8 - 76.9 | 29.0 - 69.6 |
| **Education<sup>b</sup> (years)** |  |  |  |  |  |  |
| *mean* | 14.7 | 15.3 | 14.1 | 15.5 | 15.6 | 15.4 |
| *standard deviation* | 2.0 | 2.0 | 1.9 | 2.0 | 2.3 | 1.6 |
| *range* | 12 - 19 | 12 - 19 | 12 - 18 | 12 - 19 | 12 - 20 | 12 - 18 |
| **WAIS-III Full Scale IQ** |  |  |  |  |  |  |
| *mean* | 110.3 | 110.1 | 110.5 | 111.6 | 111.2 | 112.0 |
| *standard deviation* | 15.0 | 14.2 | 16.0 | 16.3 | 19.3 | 13.1 |
| *range* | 72 - 140 | 72 - 137 | 81 - 140 | 79 - 152 | 79 - 142 | 90 - 152 |
| **WMS-III IMI** |  |  |  |  |  |  |
| *mean* | 109.7 | 114.4 | 104.7 | 111.9 | 114.8 | 109.0 |
| *standard deviation* | 16.6 | 18.3 | 13.4 | 16.9 | 16.4 | 17.4 |
| *range* | 63 - 144 | 63 - 144 | 82 - 130 | 80 - 146 | 84 - 138 | 80 - 146 |
| **WMS-III DMI** |  |  |  |  |  |  |
| *mean* | 112.6 | 116.7 | 108.3 | 111.8 | 113.5 | 110.1 |
| *standard deviation* | 17.3 | 20.4 | 12.5 | 16.0 | 14.9 | 17.2 |
| *range* | 52 - 140 | 52 - 140 | 86 - 132 | 83 - 150 | 83 - 140 | 84 - 150 |
| **Duration of Heavy Drinking<sup>cdef</sup> (years)** |  |  |  |  |  |  |
| *mean* | 17.4 | 14.3 | 20.5 | 0.0 | 0.0 | 0.0 |
| *standard deviation* | 7.7 | 5.2 | 8.5 | 0.0 | 0.0 | 0.0 |
| *range* | 5.0 - 35.0 | 6.0 - 25.0 | 5.0 - 35.0 | 0.0 - 0.0 | 0.0 - 0.0 | 0.0 - 0.0 |
| **Quantity Frequency Index<sup>cde</sup> (ounces ethanol/day; ~drinks/day)** |  |  |  |  |  |  |
| *mean* | 11.2 | 8.7 | 13.7 | 0.4 | 0.2 | 0.0 |
| *standard deviation* | 8.8 | 5.8 | 10.5 | 0.6 | 0.5 | 0.0 |
| *range* | 2.7 - 38.4 | 2.7 - 28.1 | 4.5 - 38.4 | 0.0 - 2.6 | 0.0 - 2.4 | 0.0 - 0.0 |
| **Length of Sobriety<sup>cde</sup> (years)** |  |  |  |  |  |  |
| *mean* | 8.3 | 10.6 | 5.9 | 2.1 | 3.6 | 0.5 |
| *standard deviation* | 10.3 | 11.1 | 8.8 | 6.4 | 8.5 | 1.3 |
| *range* | 0.1 - 32.3 | 0.1 - 32.1 | 0.1 - 32.3 | 0.002 - 29.2 | 0.002 - 29.2 | 0.002 - 5.1 |
| **Hamilton Rating Scale for Depression<sup>g</sup>** |  |  |  |  |  |  |
| *mean* | 3.5 | 4.9 | 2.2 | 2.4 | 3.1 | 1.8 |
| *standard deviation* | 4.2 | 4.1 | 4.0 | 2.8 | 3.3 | 2.1 |
| *range* | 0 - 18 | 0 - 17 | 0 - 18 | 0 - 12 | 0 - 12 | 0 - 8 |

Participants Characteristics ( $p < 0.05$ ): <sup>a</sup>Control Women > Control Men; <sup>b</sup>Control Men > Alcoholic Men; <sup>c</sup>Alcoholics > Controls; <sup>d</sup>Alcoholic Men > Control Men; <sup>e</sup>Alcoholic Women > Control Women; <sup>f</sup>Alcoholic Men > Alcoholic Women; <sup>g</sup>Alcoholic Women > Alcoholic Men. See Results for additional detail on number of participants.
Abbreviations: WAIS -- Wechsler Adult Intelligence Scale; WMS -- Wechsler Memory Scale; IMI -- Immediate Memory Index; DMI -- Delayed Memory Index. Five NCm and two NCw reported being lifetime abstainers.

**Supplemental Figure 1.**
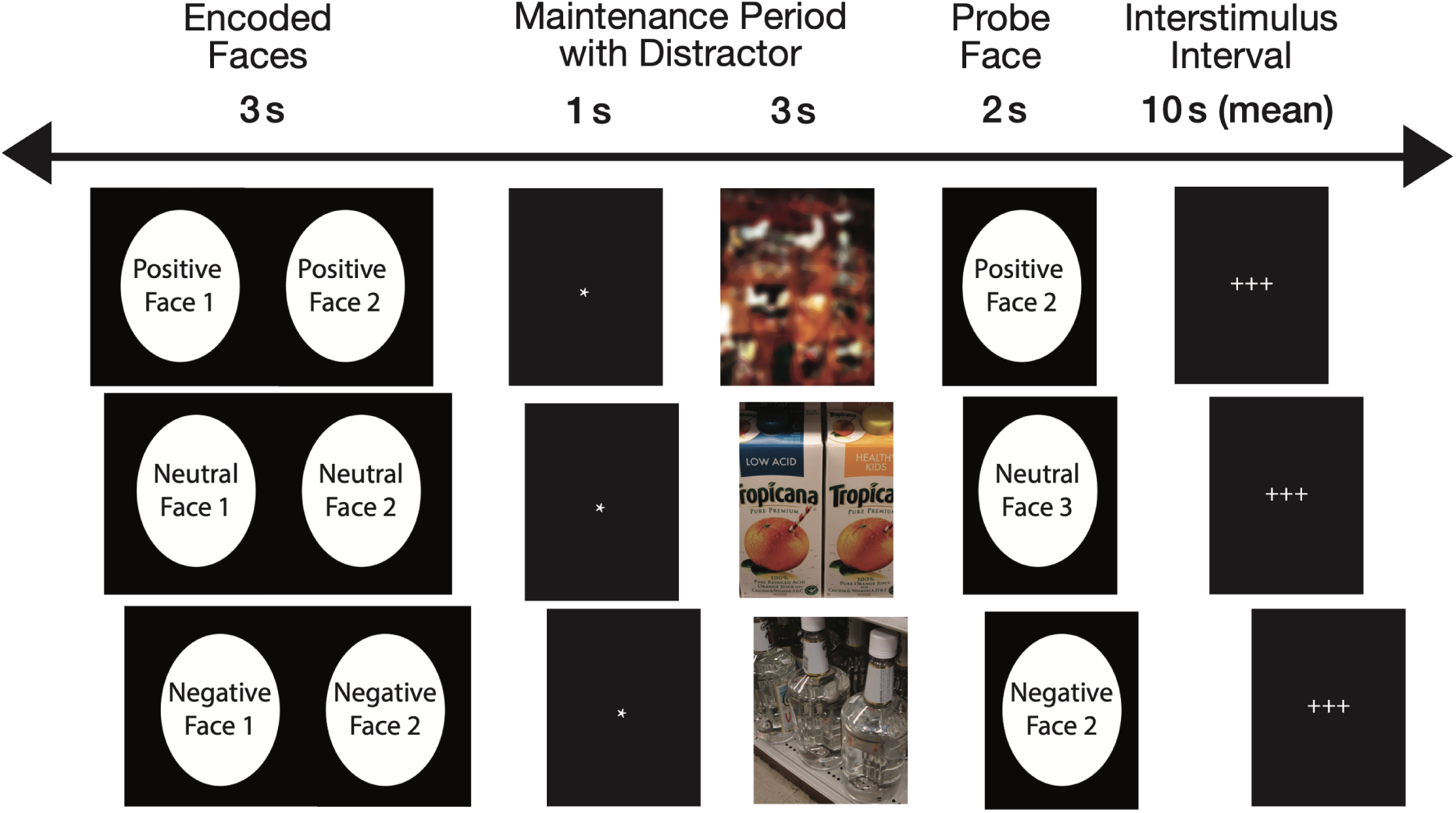
Task presented during functional neuroimaging. Two faces were presented simultaneously for 3 seconds, followed by an asterisk for one second. Next, a distractor was presented for 3 seconds. The probe face immediately followed, during which the subjects had been trained to respond with a button press with either their index or middle finger to indicate whether the probe face matched the encoded face. Three crosses served as the inter-trial interval, which lasted from 2 to 30 seconds (mean 10 seconds). While the faces in this figure have been blurred to mask the identities of the individuals, the research participants saw the original unblurred photographs.

**Supplemental Figure 2.**
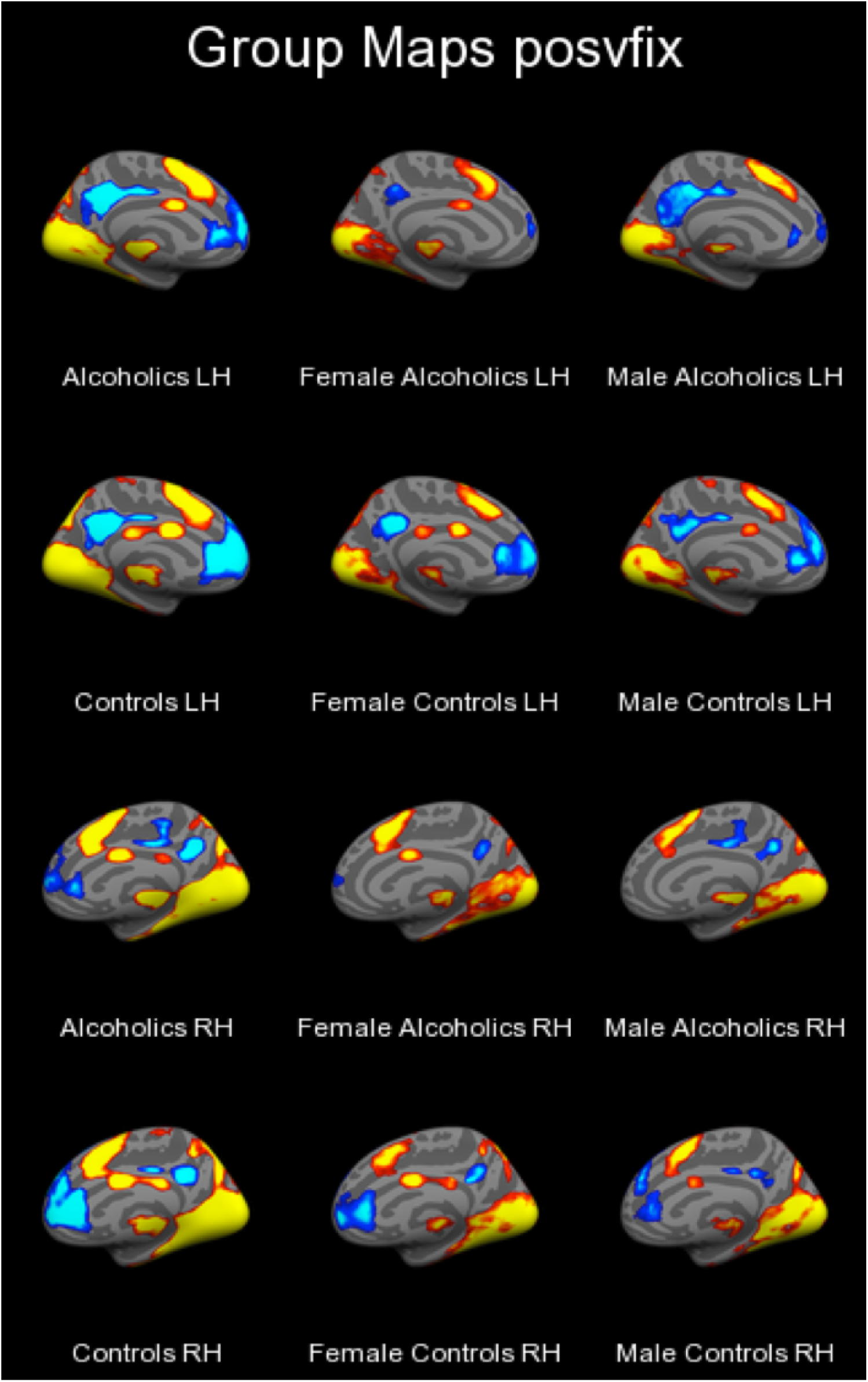
Group Cortical Surface Cluster Maps: Positive vs. Fixation, Medial. Cluster-corrected at *p*<.001 with minimum cluster size 100 mm^2^.

**Supplemental Figure 3.**
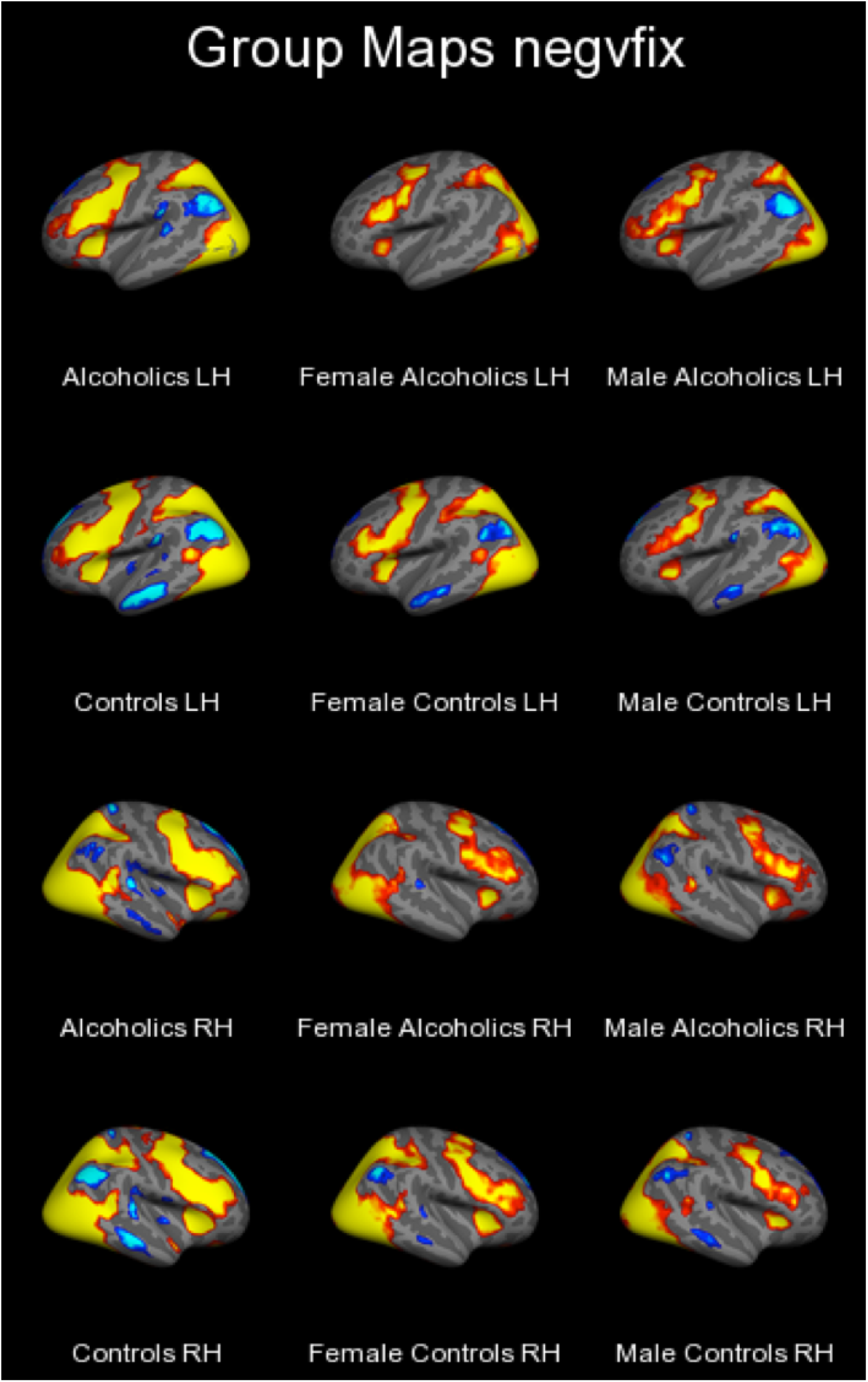
Group Cortical Surface Cluster Maps: Negative vs. Fixation, Lateral. Cluster-corrected at *p*<.001 with minimum cluster size 100 mm^2^.

**Supplemental Figure 4.**
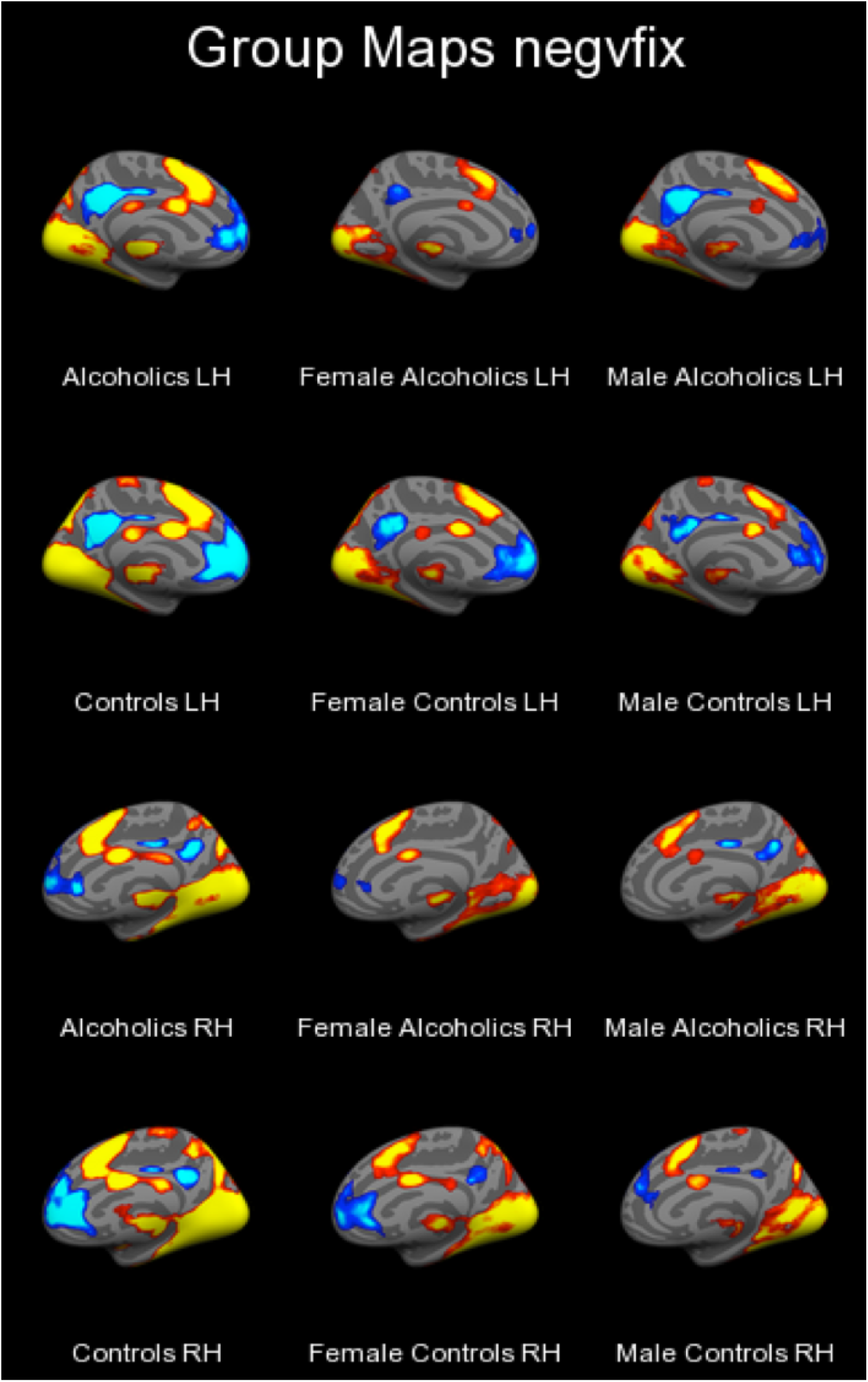
Group Cortical Surface Cluster Maps: Negative vs. Fixation, Medial. Cluster-corrected at *p*<.001 with minimum cluster size 100 mm^2^.

**Supplemental Figure 5.**
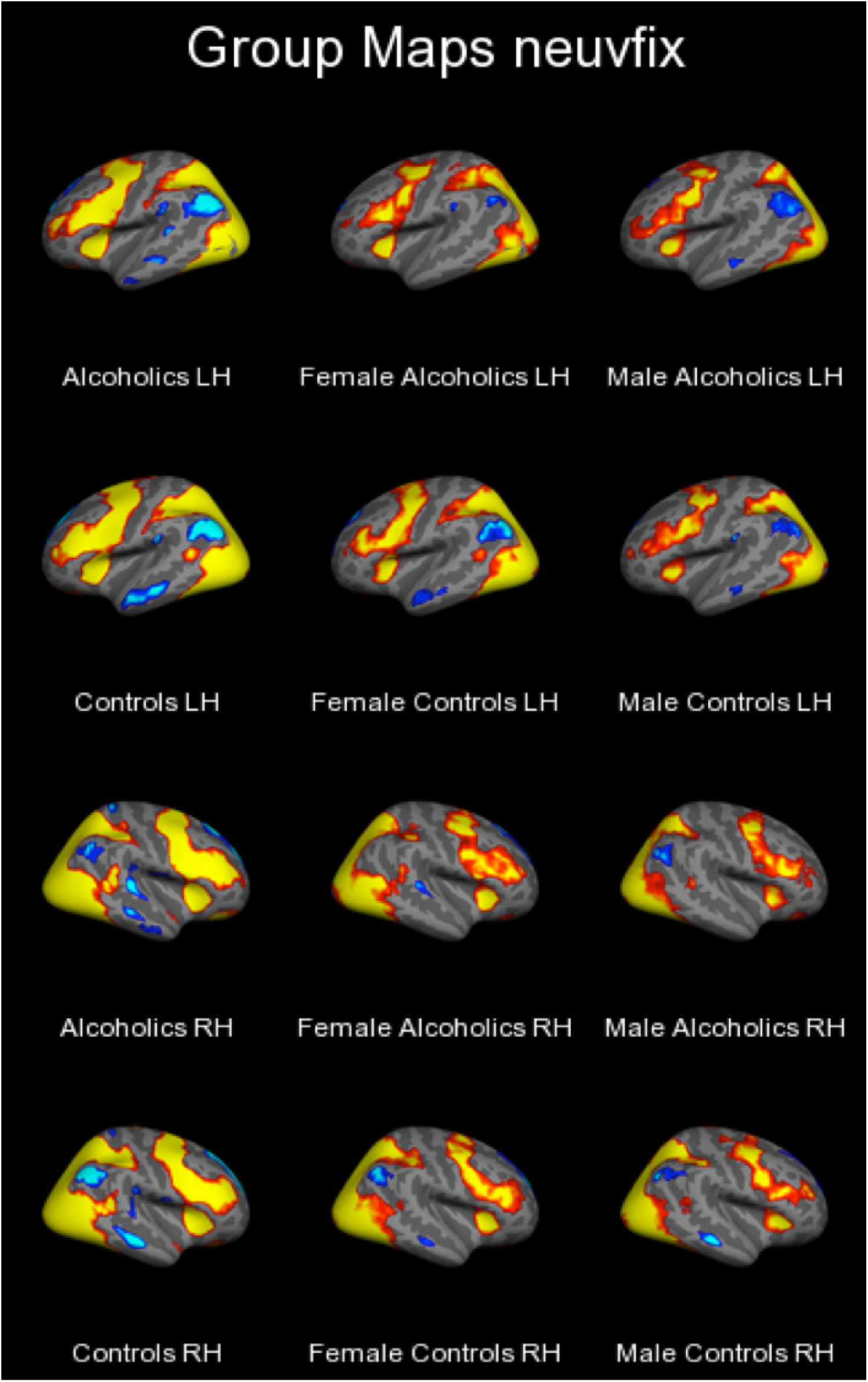
Group Cortical Surface Cluster Maps: Neutral vs. Fixation, Lateral. Cluster-corrected at *p*<.001 with minimum cluster size 100 mm^2^.

**Supplemental Figure 6.**
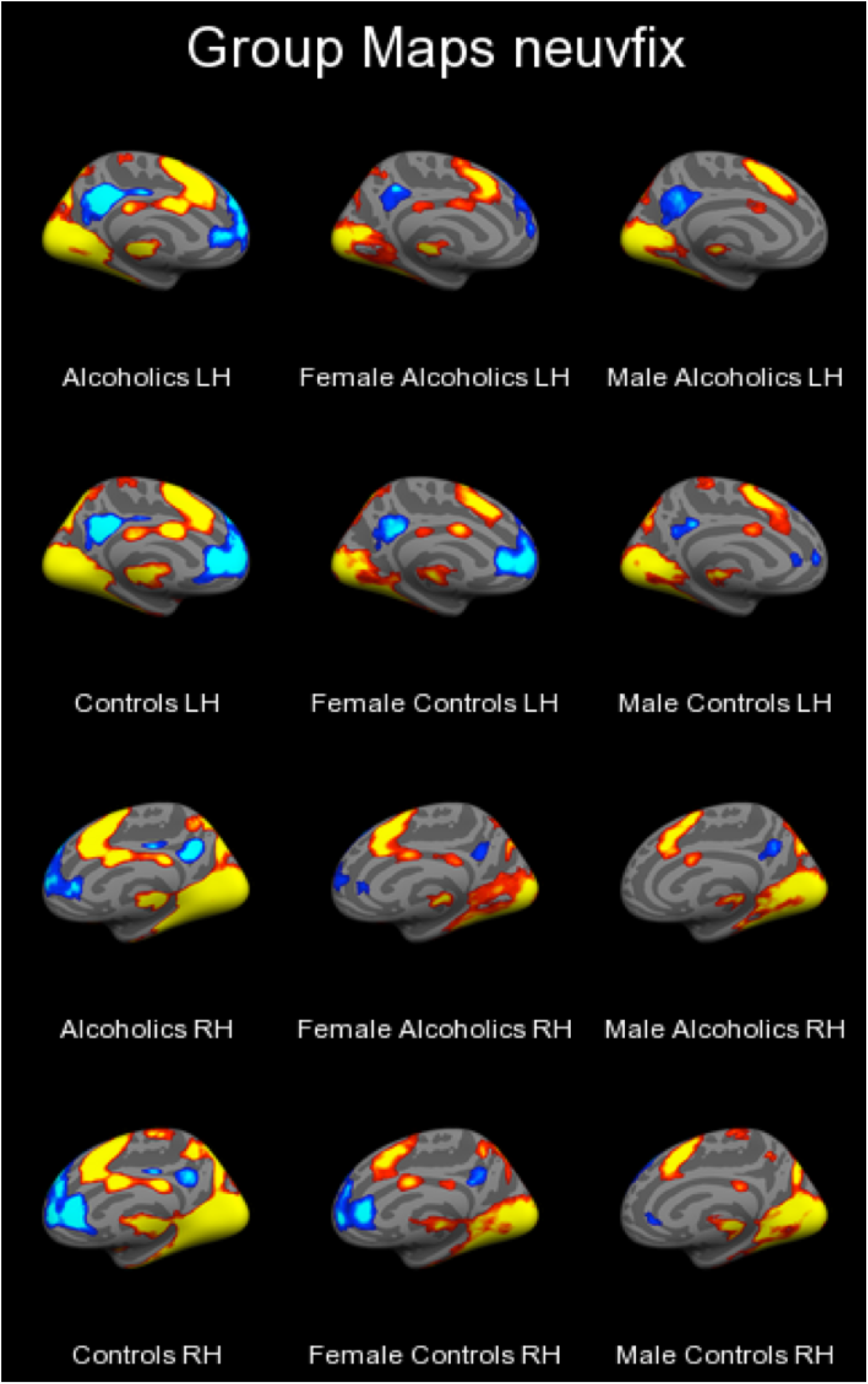
Group Cortical Surface Cluster Maps: Neutral vs. Fixation, Medial. Cluster-corrected at *p*<.001 with minimum cluster size 100 mm^2^.

**Supplemental Figure 7.**
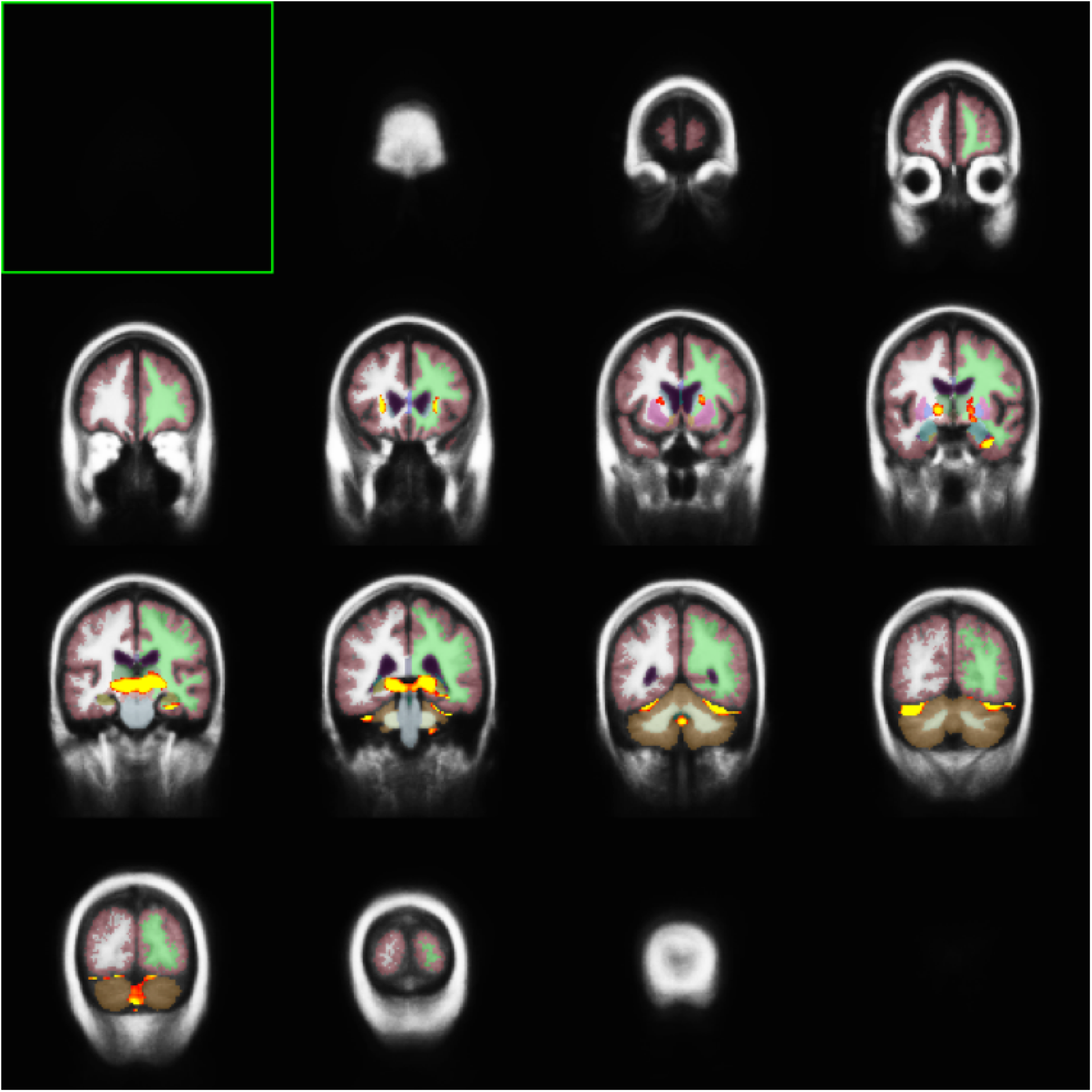
Group Subcortical Volume Cluster Maps: Alcoholic Women, Positive Faces vs. Fixation. Cluster-corrected at *p*<.001 with minimum cluster size 300 mm^3^. Shown in neurological convention.

**Supplemental Figure 8.**
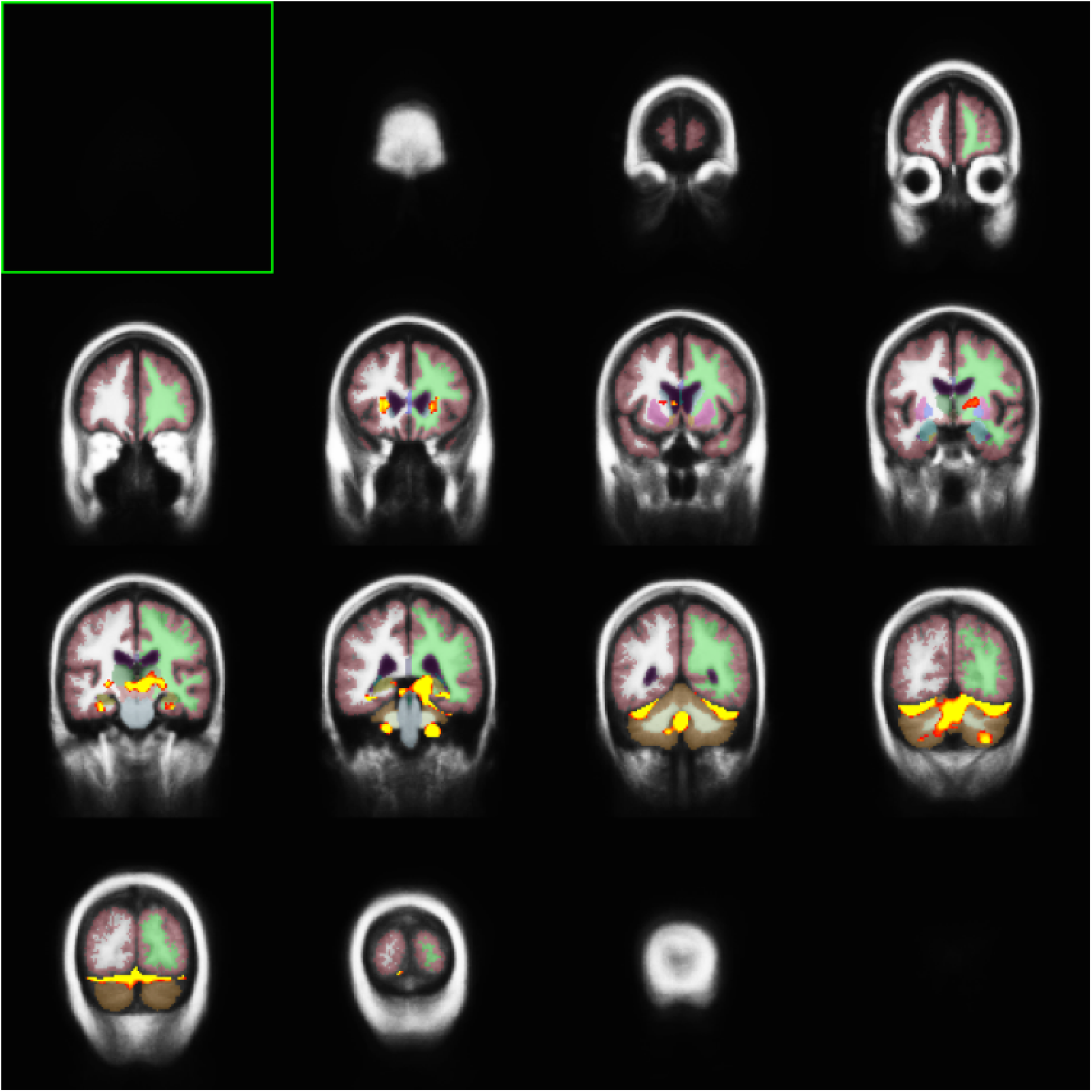
Group Subcortical Volume Cluster Maps: Alcoholic Men, Positive Faces vs. Fixation. Cluster-corrected at *p*<.001 with minimum cluster size 300 mm^3^. Shown in neurological convention.

**Supplemental Figure 9.**
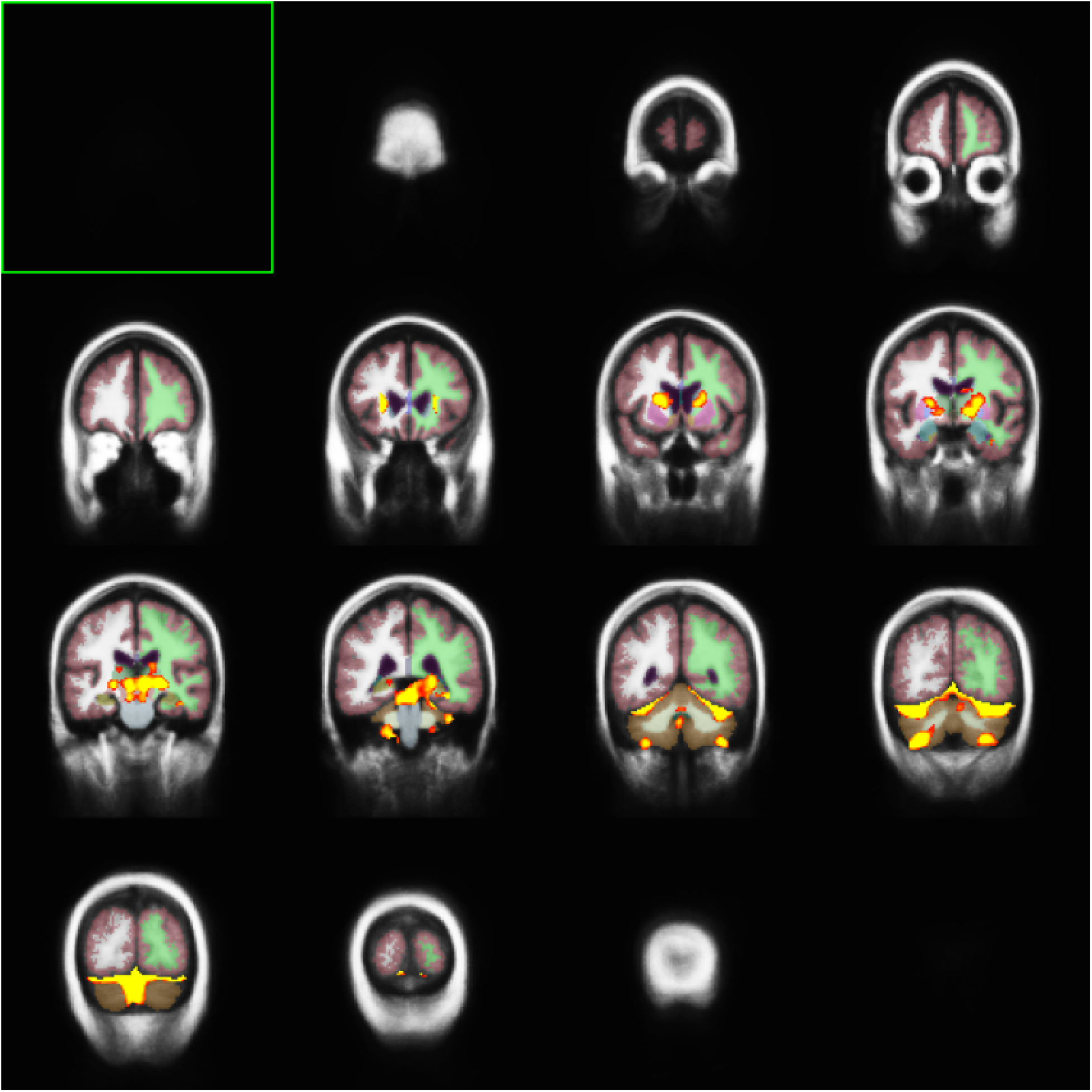
Group Subcortical Volume Cluster Maps: Control Women, Positive Faces vs. Fixation. Cluster-corrected at *p*<.001 with minimum cluster size 300 mm^3^. Shown in neurological convention.

**Supplemental Figure 10.**
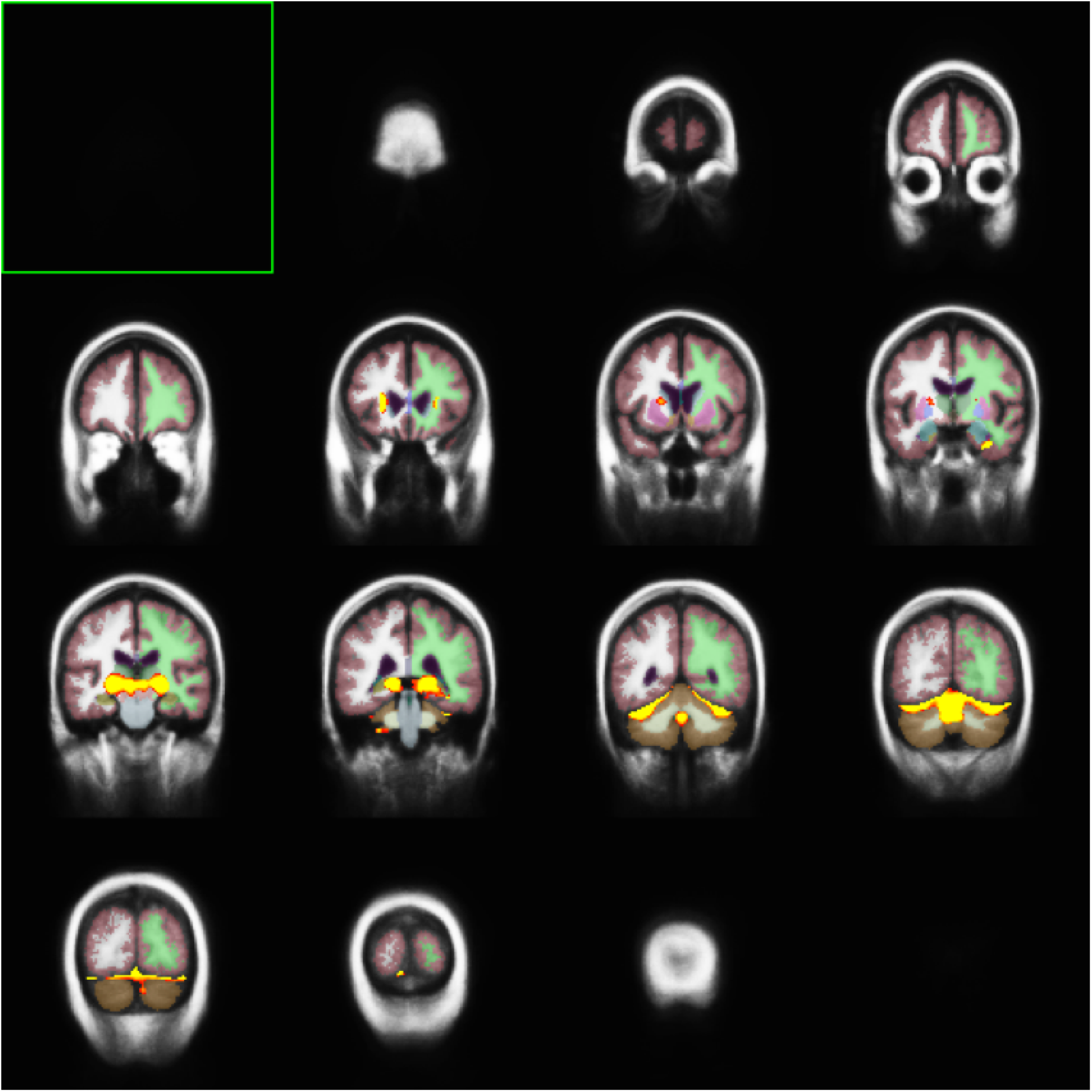
Group Subcortical Volume Cluster Maps: Control Men, Positive Faces vs. Fixation. Cluster-corrected at *p*<.001 with minimum cluster size 300 mm^3^. Shown in neurological convention.

